# Toxicological evaluation of Vanillin Flavor in E-Liquid Aerosols on Endothelial Cell Function: Findings from the Replica Project

**DOI:** 10.1101/2024.01.20.576442

**Authors:** R. Emma, A. Sun, K. Partsinevelos, S. Rust, V. Volarevic, R. Lesmana, A. Giordano, H. Goenawan, M. I. Barliana, A. Arsenijevic, N. Kastratovic, V. Markovic, B. Spasic, A. Distefano, L. Orlando, G. Carota, R. Polosa, M. Caruso, G. Li Volti

## Abstract

**Background:** There are challenges that require collaboration among researchers to ensure that tobacco harm reduction strategies are evidence-based. One key challenge is evaluating the safety of flavors used in electronic cigarettes (e-cigarettes). While many flavorings are approved as food additives or deemed “generally recognized as safe” (GRAS) for ingestion, this does not guarantee their safety when inhaled. In this context, the international research group Replica replicated a study conducted by Fetterman and colleagues in 2018, investigating the effects of aerosolized vanillin - one of the most popular flavors in vaping - on vascular endothelium when vaporized by an electronic cigarette.

**Methods:** We used Aspire Zelos 3 e-cigarette and prepared e-liquids containing propylene glycol, vegetable glycerin and vanillin. The e-liquids were vaporized under two settings - regular (1 ohm coil using wattage control mode at 14 watts) and sub-ohm (0.3 ohm coil using temperature control mode at 200 °C) – using a vaping machine, following the standardized puffing regime, ISO20768:2018. The vapor was then collected into a trapping solution to prepare aqueous extracts for the treatment of human aortic endothelial cells. We evaluated cytotoxicity, oxidative stress, nitric oxide bioavailability, and inflammation addressing some gaps reported in the original study.

**Results:** We observed some harmful effects, mostly attributable to ethanol, used to dilute vanillin in the original work by Fetterman, but no harmful effects on cell viability, their ability to produce nitric oxide, or oxidative stress from vanillin. Furthermore, no pro-inflammatory effects of vanillin were observed in terms of ICAM-1 and IL-6 gene expression.

**Conclusions:** Our results confirm the endothelial cell dysfunction observed in the original paper, but clarify that these effects are mainly attributable to ethanol and not to vaporized vanillin. These findings suggest that vanillin could be a safer flavoring agent for e-cigarette, without causing adverse effects on the cardiovascular system.

## Introduction

Electronic cigarettes (ECs) have gained increased popularity among people who smoke for their potential for smoking cessation and harm reduction^1–3^ due to their cost-effectiveness^4^ and their ability to mimic the smoking experience without the production of harmful combustion or smoke^5, 6^. Research from five independent academic institutions, involved in a broader international initiative focused on the replicability of scientific studies, further supports the reduced harm associated with ECs relative to combusted cigarettes. The consortium replicated studies examining the cytotoxic and inflammatory effects on human bronchial cells. Their findings concluded that aerosols from combustion-free nicotine delivery technologies are approximately 80% less cytotoxic compared to tobacco smoke^7^. Additionally, in a separate study, the consortium demonstrated that while tobacco combustion in cigarette smoke exhibited high levels of cytotoxicity, mutagenicity, and genotoxicity in vitro, the aerosol from ECs displayed modest or no such effects^8^. Finally, another study conducted by this consortium demonstrated a reduced impairment of repair mechanisms in vascular endothelium exposed to ECs and heated tobacco products (HTP) aerosol compared to combustion cigarette smoke^9^. Yet, despite the absence of tobacco combustion, the inhalation of these aerosols is not without potential risks.

E-cigarettes primarily consist of a battery and an atomizer where a liquid solution (commonly known as e-liquid or vape juice) is stored and vaporized by energy supplied to an electrical resistance. The liquid mainly contains propylene glycol and glycerol, with the option to include nicotine. A significant characteristic of the e-cigarette market is the availability of a variety of flavorings in e-liquids. Besides tobacco-like flavors, consumers can choose flavors such as mint, fruits, desserts, candies beverages, and many more. It is estimated that several thousand e-liquid flavors have been identified^10, 11^. These flavors are fundamental components of the vaping experience and are created using a combination of food-grade flavors, propylene glycol (PG), and vegetable glycerin (VG). Client preferences for flavors can vary significantly, making the search for the right flavor a subjective and individual experience. In the largest survey ever performed on e-cigarette use, involving almost 70,000 participants, it was found that non-tobacco flavors, especially fruit and dessert flavors, significantly contribute to successful smoking cessation among adults who formerly smoked^12^. These flavors were also considered important not only in their effort to quit smoking but also in preventing relapse to cigarette smoking.

One of the most popular and concentrated flavoring chemicals in dessert-flavored e-liquids is vanillin. Vanillin has been approved as a food additive and is considered “generally recognized as safe” (GRAS) for certain uses in food. However, this does not in itself mean that the flavorings are safe when used via inhalation^13^. The food additive approval or GRAS status of a substance only applies to specific intended uses in the food and is not supported by studies that consider inhalation toxicity. In this regard, the toxicity of flavor chemicals should be re-evaluated, particularly regarding inhalation. Concerns have been raised about specific flavoring compounds, such as diacetyl, acetyl propionyl and acetoin, which were associated with respiratory problems if inhaled ^13^. Many flavors used in e-liquids have been studied to determine their safety in vaping and tobacco products. Some studies suggest that certain flavors, including vanillin, may induce harmful effects when vaporized ^14–18^. However, many of these studies do not accurately replicate the vaporization process that occurs in e-cigarettes.

To address this, we chose to replicate the study by Fetterman et al. 2018 ^16^, using standardized methods, ISO-compliant vaping machines, and commercially available e-cigarettes to verify whether the results were reproducible and applicable to real vaping experiences. As is customary with the Replica study, these experiments were independently conducted in four international laboratories, following standardized and harmonized SOPs ^7–9^.

## Methods

### Study design and Harmonization process

This is an interlaboratory *in vitro* study conducted in the framework of the new phase of REPLICA 2.0 project. The multi-center research network established for this study includes laboratories from Italy (LAB-A) – the leading center –, USA (LAB-B), Indonesia (LAB-C), and Serbia (LAB-D). As recommended by the Centre for Open Science transparency and openness promotion guidelines (https://www.cos.io/initiatives/top-guidelines), protocols were standardized across laboratories with standard operative procedures (SOPs) defined for each experimental step. Moreover, all laboratories used the same cell line, cell-exposure equipment, and methods to assess endpoints. LAB-A arranged a kick-off meeting to train the staff of international partners and harmonize the SOPs. These SOPs have been designed to be as close as possible to the protocols of the original paper, except for the vanillin vaporization and cytotoxicity assay. Particularly, the major limitations stated by the authors of Fetterman et al.^16^ were the use of flavor diluted in ethanol and not in propylene glycol (PG) and glycerol (VG), and their heating without using an electronic cigarette and a standardized exposure system. We covered these gaps by dissolving the vanillin in PG and VG solvents and using an electronic cigarette vaped by a standardized vaping machine (LM4E; Borgwaldt; Hamburg, Germany). Another difference from the original paper is the use of neutral red uptake (NRU) assay and MTS assay instead of TUNEL for the evaluation of cytotoxicity.

### Cell culture

The experiments were performed using the Human Aortic Endothelial Cells (HAEC) purchased from Lonza (CC-2535). Cells were cultured in EBM™-2 Basal Medium (Lonza, CC-3156) supplemented with EGM™-2 SingleQuots™ (Lonza, CC-4176), and incubated in a humidified atmosphere (5% CO_2_) at 37 °C. All experiments with HAECs were performed within passages 2 - 4. When the cells reached confluence, they were detached with 0.05% trypsin–0.02% EDTA solution and replated in new flasks (cell passaging), or into 96-well plates (cell viability assay, Nitric Oxide bioavailability, oxidative stress assessment), or into 12-well plates (RT-qPCR). The cells were received by the vendor in a frozen vial, and the thawed cells have been considered as “passage 0”.

### Test products and preparation of e-liquids

The e-cigs used were Aspire Zelos 3 bought from Italian dealers. For the preparation of the e-liquids were used propylene glycol (PG) (Sigma Aldrich, P4347-500ML), vegetable glycerol (VG) (Sigma Aldrich, 1370282500), vanillin (Sigma Aldrich, V1104-100G), and absolute ethanol (Sigma Aldrich, 1009832500). The day before the exposure run were prepared 4 different liquids: PG/VG 50:50 (50% of PG and 50% of VG), PG/VG 50:50 with Vanillin (50% of PG and 50% of VG at 200 mM of vanillin), PG/VG 30:70 (30% of PG and 70% of VG), and PG/VG 30:70 with Vanillin (30% of PG and 70% of VG at 200 mM of vanillin). Vanillin was first diluted with absolute ethanol to a concentration of 2 M and then was mixed with PG and VG. PG/VG 50:50 and PG/VG 50:50 with Vanillin were used with e-cig set in Wattage control mode (1.0 Ohm coil and 14 Watt) and the MTL (mouth-to-lung) drip tip inserted (Regular setting). PG/VG 30:70 and PG/VG 30:70 with Vanillin were used with e-cig set in Temperature control mode (0.3 Ohm coil and 200 °C) and the DTL (direct-to-lung) drip tip inserted (Sub-ohm setting).

### Vapor exposure

The different e-liquids loaded to the e-cigs were vaped by the LM4E Vaping Machine (Borgwaldt, Hamburg, Germany) following the “CORESTA Reference Method n. 81” (CRM81) regimen (55 ml puff volume, drawn over 3 s, once every 30 s with square shaped profile, and with a puff velocity of 18.3 ml/s), accredited into ISO 20768:2018 ^19^. A button pre-activation of 1 s was also applied as per guidelines CRM81. The standard exhaust time for LM4E was 0.7 s with a flow rate in the impinger of 78.57 ml/s. The vapor was bubbled into an impinger filled with 30 ml of trapping solution (20% of absolute ethanol and 80% of PBS) to prepare the aqueous extracts (AqEs) for treatments. The laboratory conditions were checked using temperature and humidity sensors prior and during the exposure, maintaining a relative humidity of 40-70% and a temperature between 15 °C and 25 °C ±2°C, according to ISO 20768 ^19^.

### Neutral Red Uptake (NRU) assay

24 hours prior to AqEs treatment, HAECs were trypsinized (0.05% Trypsin/0.02% EDTA), resuspended in EGM™-2 complete medium, seeded in a 96-well plate at a density of 10 x 10^3^ viable cells/well, and incubated at 37 °C (5% CO2, humidified atmosphere). After vapor exposure, AqEs were filtered with 0.2 μm syringe filter. Then the cells were treated with seven different dilutions (1:2 or 100 mM, 1:4 or 50 mM, 1:8 or 25 mM, 1:20 or 10 mM, 1:200 or 1 mM, 1:2000 or 0.1 mM, 1:20000 or 0.01 mM) of PG/VG 50:50 AqE, PG/VG 50:50 with Vanillin AqE, PG/VG 30:70 AqE, and PG/VG 30:70 with Vanillin AqE for 24 hours. Following Fetterman’s indications ^16^, the molarity concentration reported and the relative dilutions of the AqE are referred to the initial concentration of Vanillin (200 mM) calculated in the relative liquids. For the dilution of AqEs was used EGM™-2 complete medium with 20 mM HEPES (Gibco, 15630-080). Next, the cells were washed with PBS and incubated with Neutral Red Solution (0.05 g/L; Sigma, N2889-100ML) diluted in EGM™-2 complete medium with 20 mM HEPES at 37 °C (5% CO2, humidified atmosphere) for 3 hours. Afterwards, the cells were washed with PBS, and the Neutral Red dye retained by cells was extracted by the addition of destaining solution (ethanol, distilled water, and acetic acid at a ratio of 50:49:1, respectively). Subsequently, the plates were shaken using an orbital agitator for 10 min at 300 rpm. Eventually, Neutral Red absorbance was measured in a microplate reader at 540 nm. NRU data were normalized to control medium or to vehicle control.

### MTS assay

Cell seeding and treatment procedures were performed as previously stated for the NRU assay. After 24 hours of incubation with AqEs, the treatment was removed from the wells and the cells were incubated with 100 μl of EGM™-2 complete medium with 20 mM HEPES and 20 μl of MTS (Promega, CellTiter 96® AQueous One Solution Reagent) at 37 °C (5% CO2, humidified atmosphere) for 3 hours. The absorbance of soluble formazan produced by cellular reduction of MTS was measured using a microplate reader at 490 nm. MTS data were normalized to control medium or to vehicle control.

### Oxidative Stress assessment

HAECs were seeded in a 96-well plate at a density of 10 x 10^3 cells/well and incubated at 37°C (5% CO2, humidified atmosphere) for 24 hours, until 80-90% confluence. Then the cells were exposed for 90 minutes to four different dilutions (1:20 or 10 mM, 1:200 or 1 mM, 1:2000 or 0.1 mM, and 1:20000 or 0.01 mM) of PG/VG 50:50 AqE, PG/VG 50:50 with Vanillin AqE, PG/VG 30:70 AqE, and PG/VG 30:70 with Vanillin AqE. The reported molarity concentration and dilutions of AqEs were established basing on the initial Vanillin concentration of the related products (200 mM). Prior to dilution, AqEs were filtered with 0.2 μm syringe filter. For the dilution was used EGM™-2 complete medium with 20 mM HEPES. Next, HAECs were incubated for 30 minutes with 10 μmol/L dihydroethidium (DHE; Thermo Fisher Scientific, D11347). After that, HAECs were washed 3 times with pre-warmed DPBS to remove DHE. As a positive control was used 50 μmol/L Antimycin A (Sigma-Aldrich, A8674) for 30 minutes. The fluorescence intensity was measured within 30 minutes using a microplate reader with excitation of 518 nm and emission of 606 nm. Data are shown as fold change in DHE fluorescence in comparison with vehicle control.

### Nitric Oxide Bioavailability

Cell seeding and exposure to AqE solutions were performed as explained above (Oxidative Stress assessment section). Next, the cells were incubated for 30 minutes with 3 µmol/L 4,5- diaminofluorescein diacetate (DAF-2 DA; Sigma-Aldrich, D225). After that, the cells were washed 2 times with pre-warmed DPBS, stimulated for 15 minutes with 1 µmol/L Calcium Ionophore A23187 (Sigma-Aldrich, C7522), and fixed with 2% paraformaldehyde in DPBS for 10 minutes at room temperature. Afterwards, paraformaldehyde was replaced with 200 μl of DPBS to each well and the fluorescence intensity (excitation of 492 nm, emission peak at 515 nm) was measured by a microplate reader. Nitric Oxide data were presented as percentage increase in DAF-2 DA fluorescence stimulated by Calcium Ionophore A23187 compared with unstimulated cells.

### Gene expression

HAECs were seeded in 12-well plates at a density of 100 x 10^3^ cells/well and incubated at 37 °C (5% CO2, humidified atmosphere) for 24 hours, until 80-90% confluence. Then the cells were exposed to 1:20, 1:200, 1:2000, and 1:20000 dilutions of AqEs. AqE solutions were prepared as previously described for oxidative stress assessment. Next, the cells were incubated for 90 minutes with complete medium (20 mM HEPES) to allow changes in RNA expression. For cell disruption and RNA isolation was used the RNeasy Mini Kit (Qiagen) following the instructions provided by the manufacturer. RNA purity and quantification were performed using spectrophotometric absorbance measurements at 260 nm and 280 nm. cDNA synthesis of the isolated RNA was carried out with the High-Capacity cDNA Reverse Transcription Kit (Thermo Fisher Scientific) according to the manufacturer’s protocol. Quantitative Real-Time PCR was performed using the TaqMan™ Fast Advanced Master Mix (Thermo Fisher Scientific) following the manufacturer’s instruction manual. As reference gene was used glyceraldehyde 3-phosphate dehydrogenase (GAPDH). GAPDH, IL-6, and ICAM-1 TaqMan™ probes were purchased from Thermo Fisher Scientific. The 2^−ΔΔCt^ was calculated from cycle threshold (Ct) values, after normalization to GAPDH as housekeeping gene.

### Statistics

Microsoft Excel was used to tabulate and process all of the raw data. The Shapiro-Wilk test was used for assessing the normality or skewness of data distribution. Correlation analyses were performed to evaluate the relationship between the results of each laboratory. Pearson’s correlation analysis was conducted for symmetrical data, while Spearman’s Rank correlation analysis was used for skewed data. Moreover, the intra-class correlation coefficient (ICC) was computed using an absolute-agreement, two-way mixed-effects model in order to evaluate the agreement in the repeatability of the intrasession measurements among the laboratory results. R version 4.2.3 (2023-03-15) was utilized for reproducibility analyses, including the generation of correlation plots. The outlier detection was made using the robust regression-based outlier rejection (ROUT) test. All data were reported as median (Interquartile range – IQR). The Kruskal-Wallis test was applied to determine any statistically significant differences between the medians of study groups. Additionally, post-hoc Dunn’s test was performed for a more detailed examination of group differences. All analyses were considered significant with a p value of less than 5 %. GraphPad Prism 8 software was used for data analysis and generation of graphs unless otherwise stated.

## Results

### Interlaboratory reproducibility

The results of the Intraclass Correlation Coefficient (ICC) calculations using an absolute-agreement, two-way mixed-effects model are presented in Table 1.

**Table 1.**
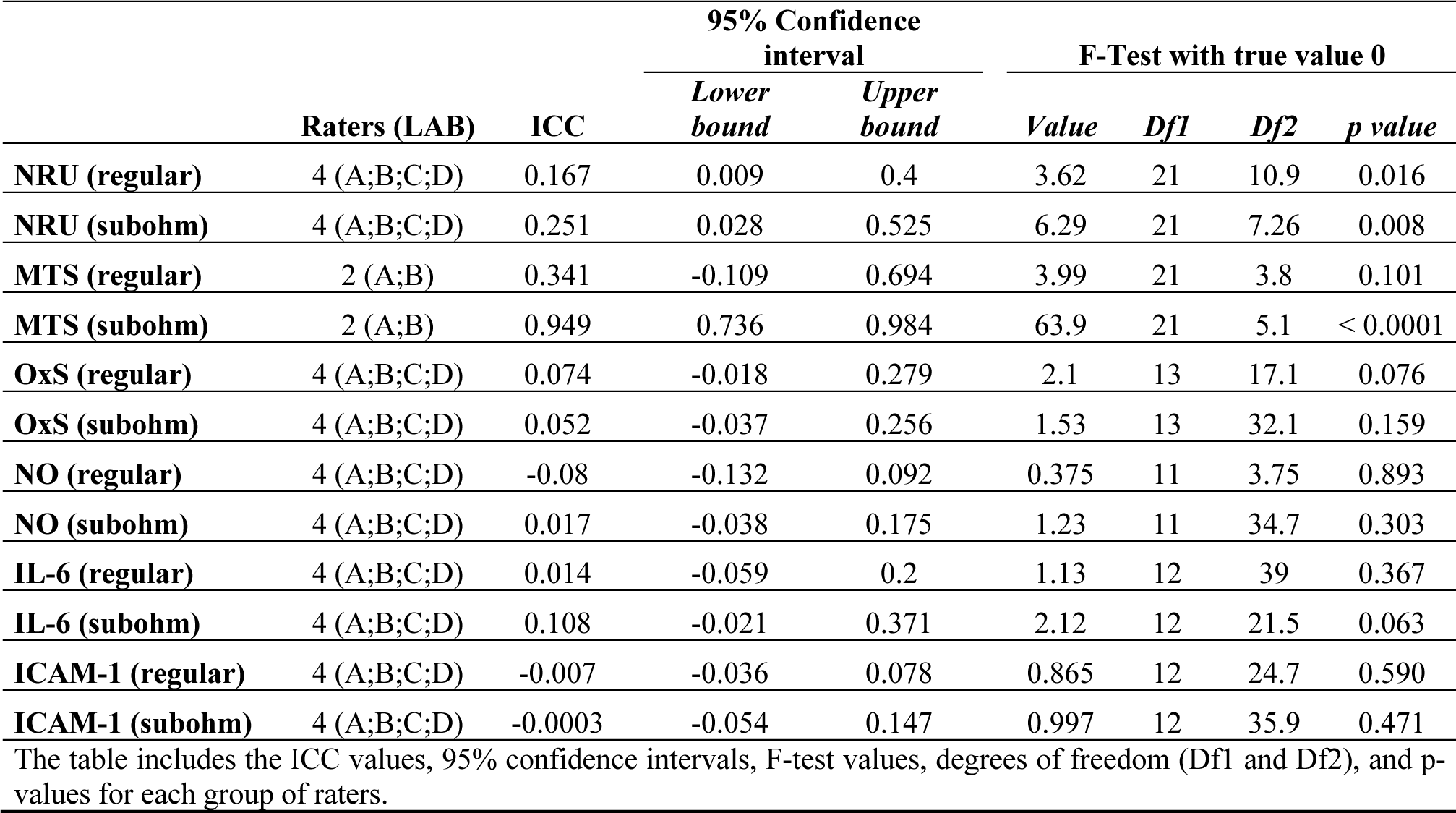
Results of ICC Calculation using absolute-agreement, two-way mixed-effects model.

The correlations of NRU results among all laboratories are shown in Figure S1 of supplementary material. Significant correlations were observed among all laboratories for the regular setting, except for the correlation between LAB-A and LAB-D. Strong correlations were also observed for the sub-ohm setting across all laboratories. Also, the NRU for regular setting had an ICC of 0.167, which was statistically significant (p= 0.016). Similarly, the NRU (sub-Ohm) group had an ICC of 0.251, also significant (p= 0.008).

For the MTS (regular) group, while correlation indicates that there is a good relationship (rho= 0.483, p= 0.024) between the measurements of the two centers (Fig. S2A), the nonsignificant ICC of 0.341 (p= 0.101) indicates that this relationship is not strong enough to ensure good reproducibility. In contrast, the MTS (sub-Ohm) group showed high correlation coefficient (rho= 0.896) (Fig. S2B) and high ICC of 0.949, indicating highly significant results (p< 0.0001).

Poor correlation was observed for Oxidative stress, Nitric oxide, IL-6 and ICAM-1 assessments for both regular and sub-ohm setting, as showed respectively in Figure S3, S4, S5 and S6 of supplementary material. Also, the ICC values indicated poor agreement among laboratories when performing oxidative stress and nitric oxide evaluations. Indeed, the oxidative stress (regular) group showed an ICC of 0.074, which was not significant (p= 0.076). The oxidative stress (sub-Ohm) group had an ICC of 0.052, also not significant (p= 0.159). For the NO (regular) group, the ICC was -0.08, which was not significant (p= 0.893). The NO (sub-Ohm) group had a low ICC of 0.017, which was also not significant (p= 0.303). For the IL-6 (regular) group, the ICC was 0.014, which was not statistically significant (p = 0.367). The IL-6 (sub-Ohm) group had a slightly higher ICC of 0.108, which approached significance (p = 0.063). For the ICAM-1 (regular) group, the ICC was -0.007, which was not significant (p = 0.590). Similarly, the ICAM-1 (sub-Ohm) group had a very low ICC of -0.0003, also not significant (p = 0.471).

These results indicate that the reliability of the measurements varied across the different evaluations and settings, with the NRU (both regular and sub-ohm) and MTS (sub-ohm) assays showing the highest reliability.

### Cytotoxicity evaluation of Vanillin

Cytotoxicity of HAECs exposed to PG/VG Vanillin for both regular and sub-ohm settings was evaluated by NRU and MTS assays after 24 hours of treatments, in contrast to Fetterman’s assessment at 90 minutes. The evaluation of cytotoxicity induced with regular setting by NRU showed significant decrease of HAECs viability after exposure to 100 mM of Vehicle control (p= 0.025), PG/VG base (p= 0.016) and PG/VG Vanillin (p= 0.012) compared to control medium. No significant differences were observed between each concentration of both PG/VG base and PG/VG Vanillin and the corresponding vehicle control concentration (Fig 1-A). NRU data normalized to vehicle control (Fig 1-B) revealed no significant differences in cell viability among PG/VG base and PG/VG Vanillin compared at each concentration (Fig 1-B).

**Figure 1.**
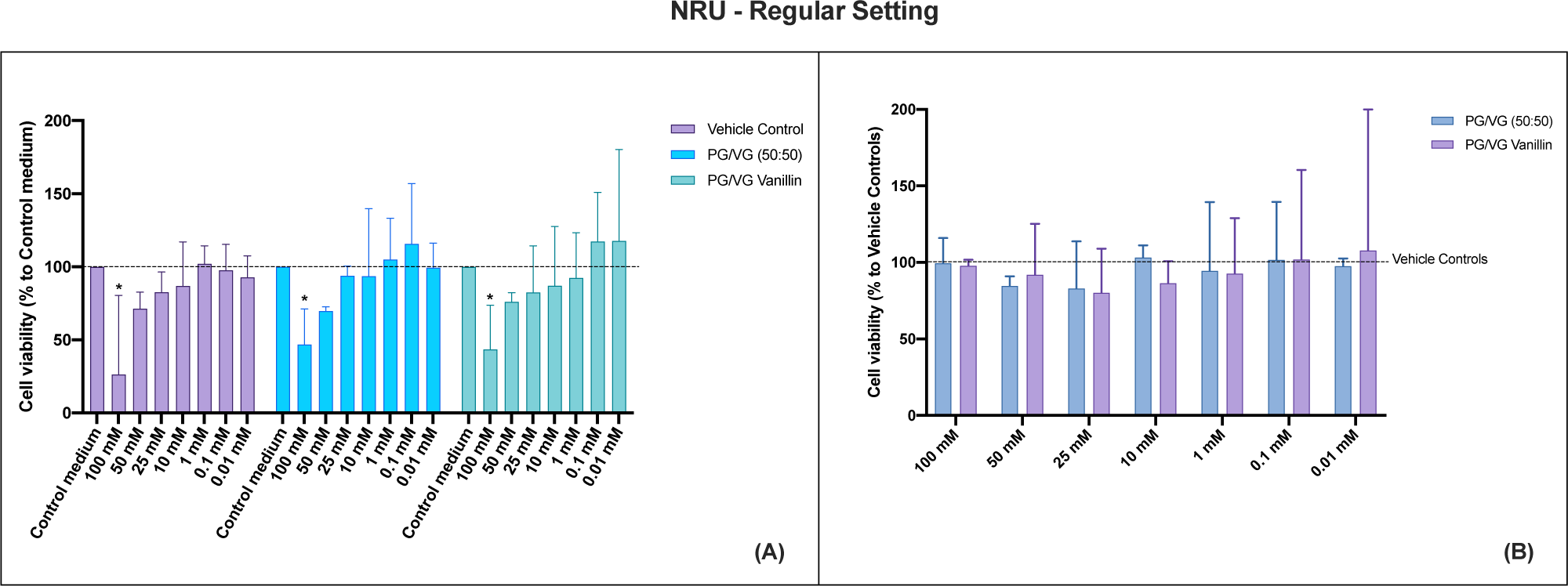
HAECs cytotoxicity evaluation by NRU assay of PG/VG base and PG/VG Vanillin using the regular setting at 24 hours. (**A**) NRU data normalized as percentage of control medium. The dashed line corresponds to control medium. (**B**) NRU data normalized as percentage of corresponding Vehicle control concentration. The dashed line corresponds to Vehicle control. All data are reported as median (IQR). * p< 0.05.

The evaluation of cytotoxicity induced with sub-ohm setting assessed by NRU assay demonstrated significant decreases of HAECs viability not only after exposure to 100 mM of Vehicle control (p= 0.0001), PG/VG base (p= 0.0002) and PG/VG Vanillin (p< 0.0001) but also to 50 mM of Vehicle control (p= 0.0032), PG/VG base (p= 0.004) and PG/VG Vanillin (p= 0.0007), and 25 mM of PG/VG Vanillin (p= 0.0236) compared to control medium. (Fig 2-A). The NRU data normalization to vehicle control (Fig 1-B) indicated no significant differences in cell viability among PG/VG base and PG/VG Vanillin compared at each concentration (Fig 2-B).

**Figure 2.**
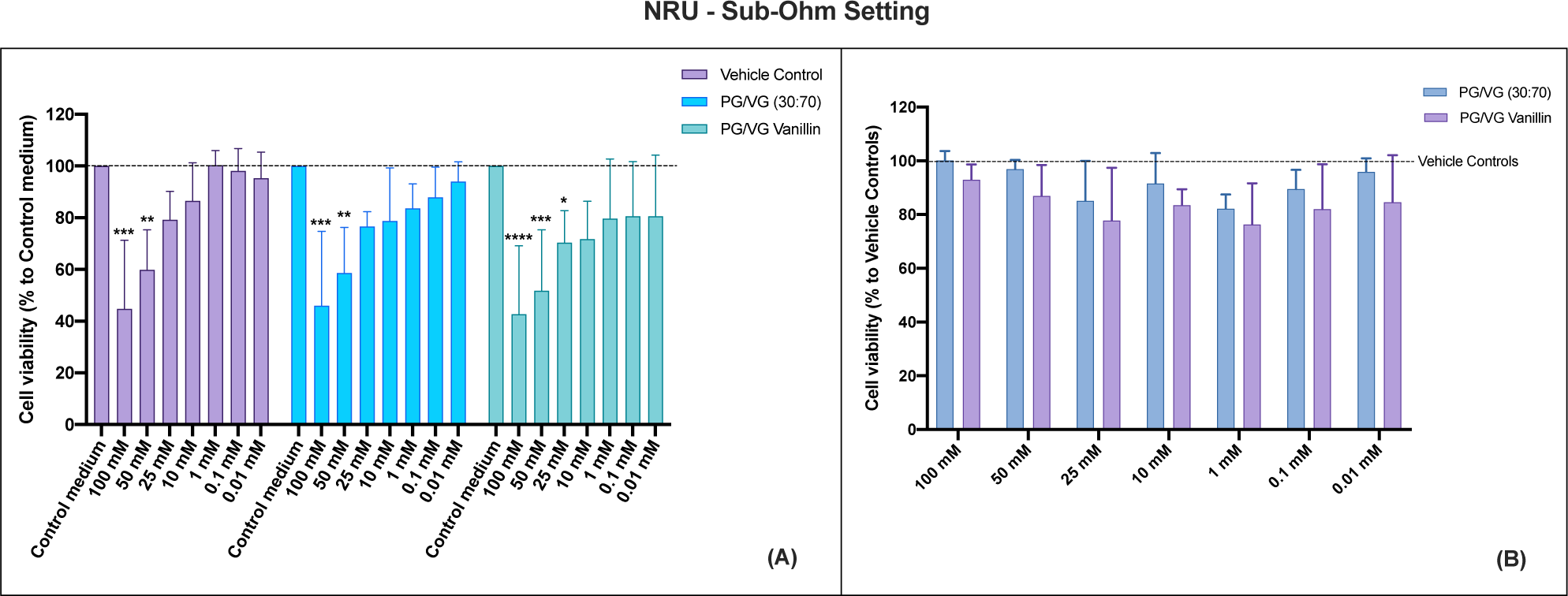
HAECs cytotoxicity evaluation by NRU assay of PG/VG base and PG/VG Vanillin using the sub-ohm setting at 24 hours. (**A**) NRU data normalized as percentage of control medium. The dashed line corresponds to control medium. (**B**) NRU data normalized as percentage of corresponding Vehicle control concentration. The dashed line corresponds to Vehicle control. All data are reported as median (IQR). *p< 0.05; ** p< 0.01; ***p< 0.001; ****p< 0.0001.

Similar results were observed when the MTS assay was used for the cytotoxicity evaluation in HAECs exposed to PG/VG and PG/VG Vanillin. The MTS assay was carried out only by LAB-A and LAB-B, thus having two independent replicants of each experiment. The assessment of cytotoxicity induced under regular setting, as determined by the MTS assay, demonstrated a reduction in the viability of HAECs following exposure to 100 mM concentrations of Vehicle control (p< 0.0001), PG/VG base (p< 0.0001), and PG/VG Vanillin (p< 0.0001) in comparison to the control medium. The 50 mM concentration of Vehicle control (p= 0.015), PG/VG base (p= 0.016), and PG/VG Vanillin (p p= 0.044) induced also a slight decrease compared to control medium. No significant variations were noted between each concentration of both PG/VG base and PG/VG Vanillin and their respective vehicle control concentrations (Fig 3-A). Normalizing the MTS data to the vehicle control (Fig 3-B) revealed no significant differences among the test products and their corresponding vehicle controls. Additionally, no notable differences in cell viability were observed when comparing PV/VG base and PG/VG Vanillin at each concentration (Fig 3-B).

**Figure 3.**
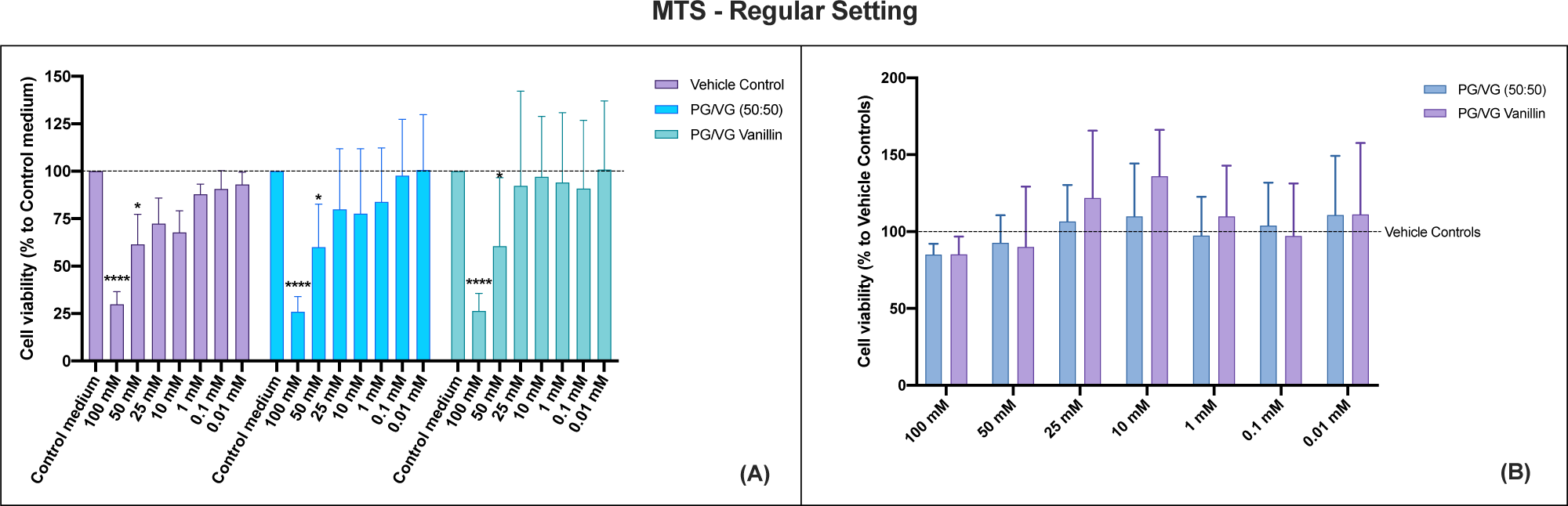
HAECs cytotoxicity evaluation by MTS assay of PG/VG base and PG/VG Vanillin using the regular setting at 24 hours. (**A**) MTS data normalized as percentage of control medium. The dashed line corresponds to control medium. (**B**) MTS data normalized as percentage of corresponding Vehicle control concentration. The dashed line corresponds to Vehicle control. All data are reported as median (IQR). *p< 0.05; **** p< 0.0001.

**Figure 4.**
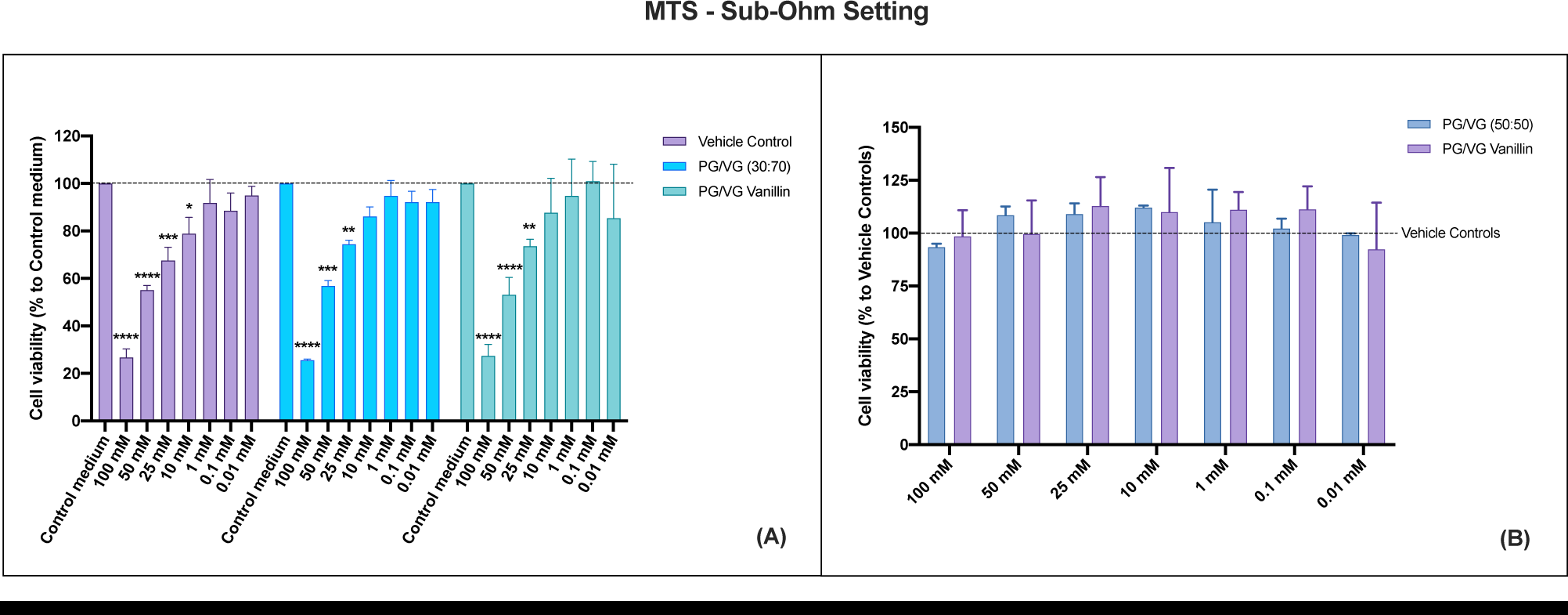
HAECs cytotoxicity evaluation by MTS assay of PG/VG base and PG/VG Vanillin using the sub-ohm setting at 24 hours. (**A**) MTS data normalized as percentage of control medium. The dashed line corresponds to control medium. (**B**) MTS data normalized as percentage of corresponding Vehicle control concentration. The dashed line corresponds to Vehicle control. All data are reported as median (IQR). *p< 0.05; ** p< 0.01; ***p< 0.001; ****p< 0.0001.

The assessment of cytotoxicity induced through sub-ohm settings by the MTS assay revealed more significant reductions in the viability of HAECs. The decrease was evident not only after exposure to 100 mM concentrations of Vehicle control (p< 0.0001), PG/VG base (p< 0.0001), and PG/VG Vanillin (p< 0.0001) but also with 50 mM concentrations of Vehicle control (p< 0.0001), PG/VG base (p= 0.0001), and PG/VG Vanillin (p< 0.0001), 25 mM of Vehicle control (p= 0.0007) PG/VG base (p= 0.002) and PG/VG Vanillin (p= 0.003), and 10 mM of Vehicle control (p= 0.014) compared to the control medium (Fig 2-A). Normalizing the MTS data to the vehicle control (Fig 1-B) revealed no significant differences among the test products and their respective vehicle controls. Furthermore, the comparison between PG/VG base and PG/VG Vanillin at each concentration indicated no significant variations in cell viability (Fig 1-B).

All these results demonstrate that the cytotoxic effect on HAECs is predominantly due to the ethanol present in the vehicle control. In fact, normalization of the data to the respective ethanol concentrations reveals no cytotoxic effect of the aerosols with and without vanillin.

### Oxidative Stress

Following 90 minutes HAECs’ treatment with PG/VG and PG/VG Vanillin at various doses, oxidative damage was measured using the fluorescent dye DHE, as reported by Fetterman and colleagues ^16^. The assessment of oxidative stress for the regular setting revealed no significant difference between the different conditions tested in comparison to the medium control (Fig. 5A). While there seems to be an increase in oxidative stress response following exposure to the vehicle control and unflavored PG/VG and PG/VG Vanillin, the considerable variability observed in the results likely precludes the statistical demonstration. Similar results were observed assessing the sub-ohm setting (Fig. 6A). No significant differences in oxidative stress between the control group and any of the experimental groups, including those treated with various concentrations of Vehicle control, PG/VG, and PG/VG Vanillin. However, the high variability, as indicated by the large interquartile ranges, points to a broad range of individual responses within each group. This variability suggests that while the overall group differences are not statistically significant, individual responses to the treatments can vary widely. Moreover, the data normalization to the corresponding Vehicle control concentrations (Fig. 5B and 6B) indicated no significant differences between unflavored PG/VG and PG/VG Vanillin compared to the respective Vehicle control concentration.

**Figure 5.**
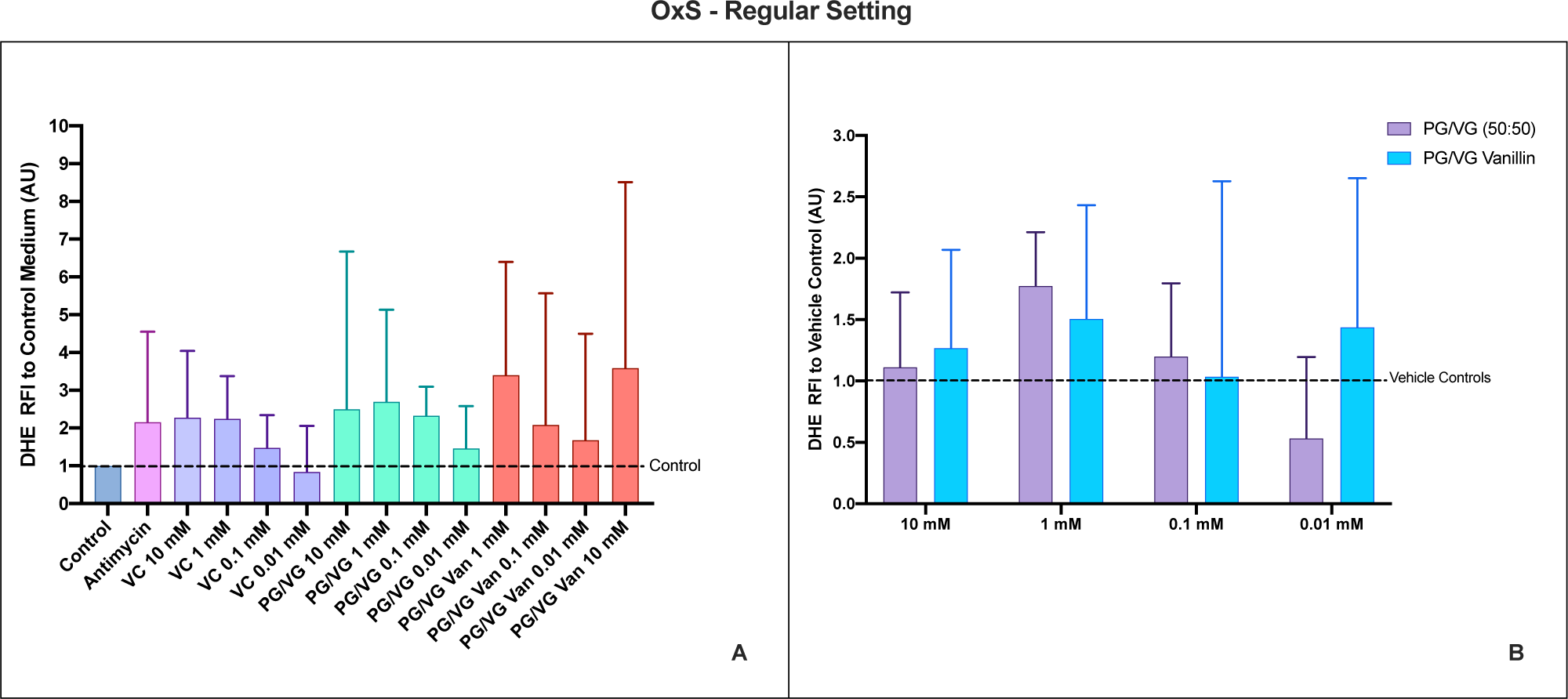
HAECs Oxidative Stress (OxS) evaluation by DHE of PG/VG base and PG/VG Vanillin using the regular setting. (**A**) OxS data normalized to control medium. The dashed line corresponds to control medium. (**B**) OxS data normalized to the corresponding Vehicle control concentration. The dashed line corresponds to Vehicle control. All data are reported as median (IQR). VC: Vehicle control

**Figure 6.**
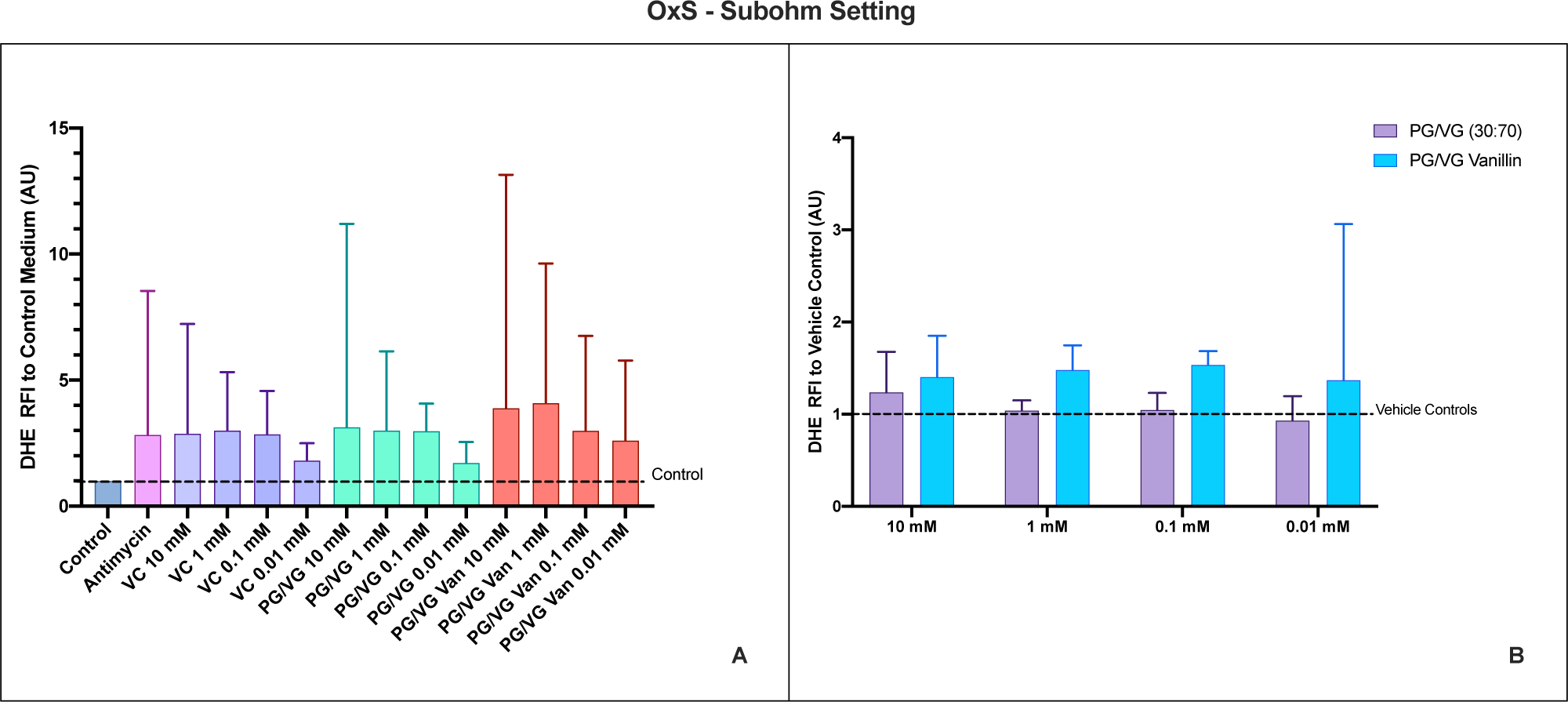
HAECs Oxidative Stress (OxS) evaluation by DHE of PG/VG base and PG/VG Vanillin using the regular setting. (**A**) OxS data normalized to control medium. The dashed line corresponds to control medium. (**B**) OxS data normalized to the corresponding Vehicle control concentration. The dashed line corresponds to Vehicle control. All data are reported as median (IQR). VC: Vehicle control

### Nitric Oxide Bioavailability was not impaired by Vanillin

As carried out by Fetterman and colleagues ^16^, we evaluated the effect of vanillin on nitric oxide production by HAECs in response to A23187 stimulation. No significant differences were observed between PG/VG and PG/VG Vanillin and the respective Vehicle Control concentrations for both regular (Fig. 7) and sub-ohm settings (Fig. 8). In some cases, it appears that only PG/VG exposure may exert a minor impact on NO production, though this is not significant compared to vehicle control. Instead, the exposure to PG/VG Vanillin appears to improve the bioavailability of NO compared to PG/VG, although no significant differences were observed among PG/VG and PG/VG Vanillin.

**Figure 7.**
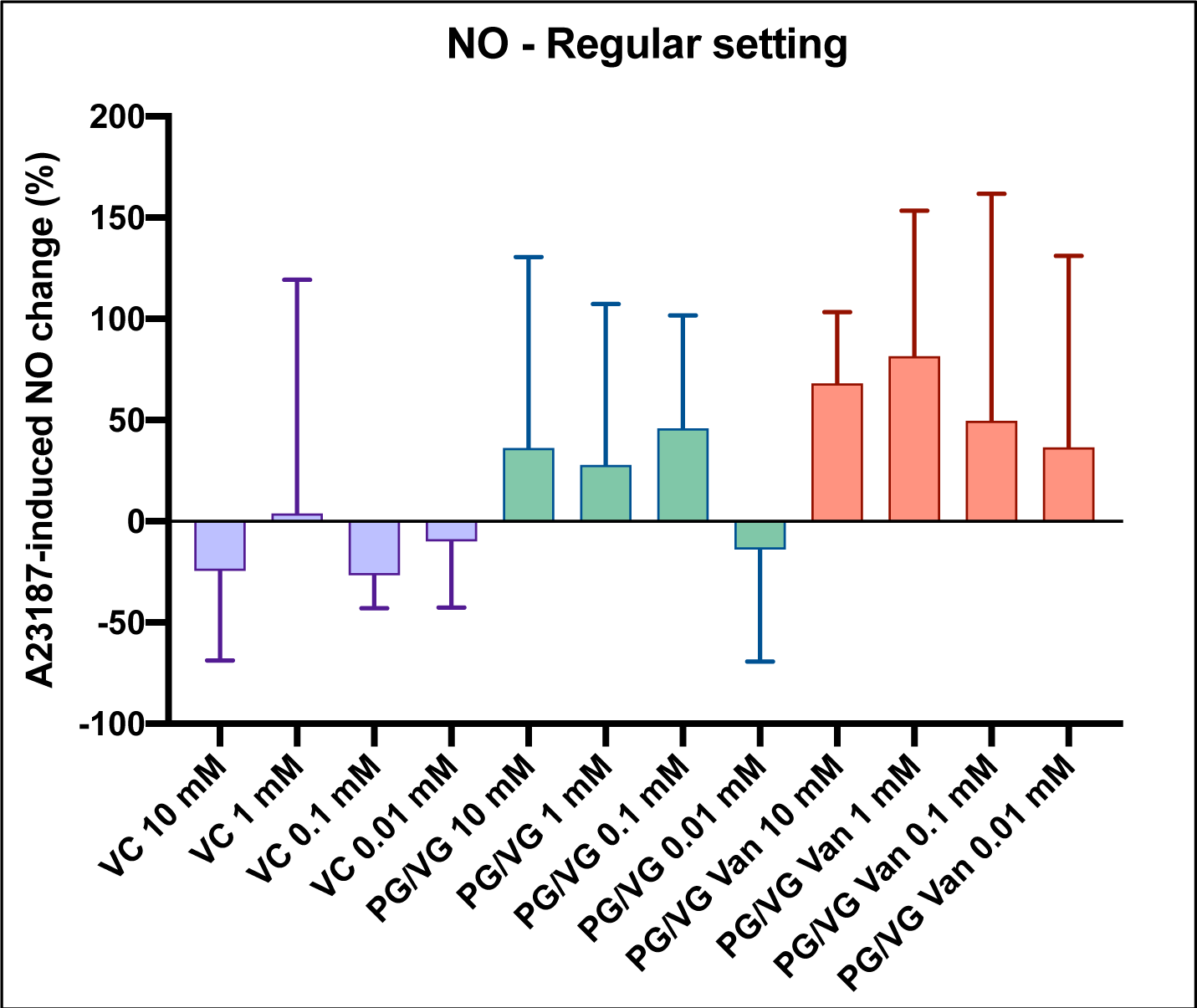
Effects of PG/VG and PG/VG Vanillin on A23187-stimulated nitric oxide production in human aortic endothelial cells (HAECs) using Regular Setting. All data are normalized as percentage of A23187-induced NO change and reported as median (IQR). VC: Vehicle control

**Figure 8.**
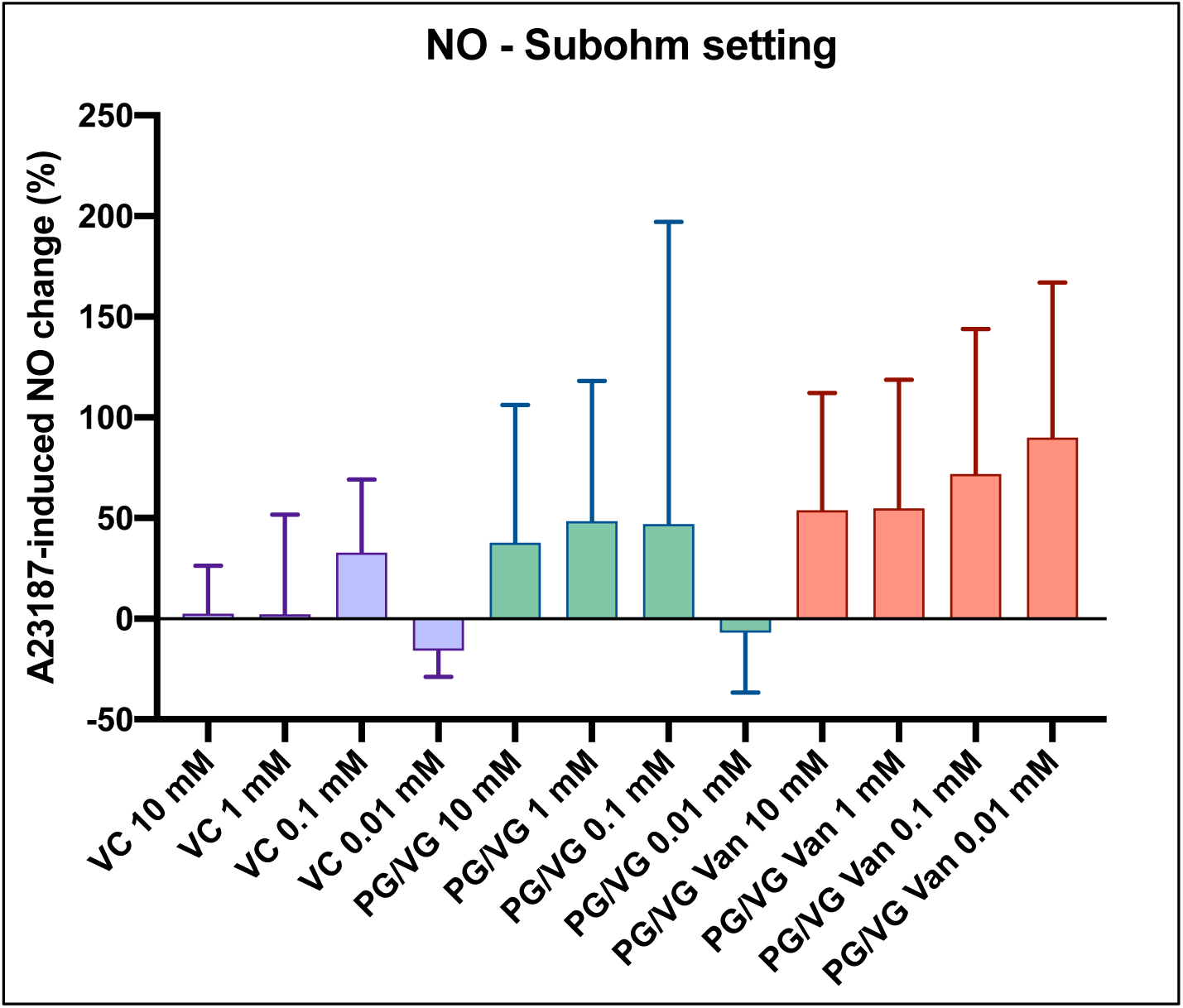
Effects of PG/VG and PG/VG Vanillin on A23187-stimulated nitric oxide production in human aortic endothelial cells (HAECs) using Sub-Ohm Setting. All data are normalized as percentage of A23187-induced NO change and reported as median (IQR). VC: Vehicle control

### Gene expression of IL-6 and ICAM-1

The evaluation of IL-6 gene expression in aortic endothelial cells treated under the experimental conditions showed no statistically significant variations compared to the control. Both PG/VG and PG/VG Vanillin treatments did not show significant differences in IL-6 expression compared to the control. This result was observed for both the Regular (Fig. 9A) and Sub-ohm (Fig. 10A) experimental setups. Similarly, the gene expression of ICAM-1 did not show statistically significant variations in aortic endothelial cells treated with Vehicle Control, PG/VG, and PG/VG Vanillin at the various concentrations (10 mM, 1 mM, 0.1 mM, and 0.01 mM). Again, no significant differences in ICAM- 1 expression were observed compared to the control in both the Regular (Fig. 9B) and Sub-ohm (Fig. 10B) setups.

**Figure 9.**
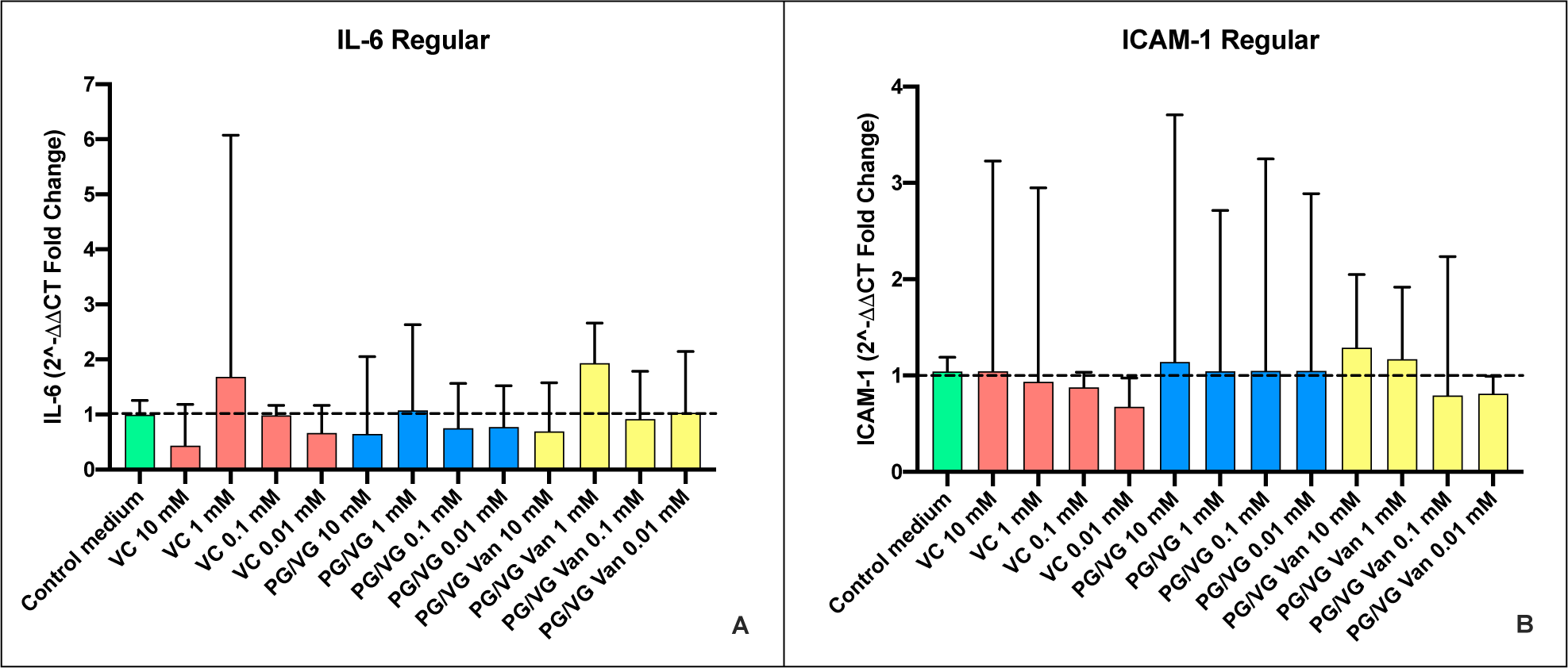
Gene expression of IL-6 (**A**) and ICAM-1 (**B**) in aortic endothelial cells under different treatments produced using the Regular setting. All data are reported as median (IQR). VC: Vehicle control.

**Figure 10.**
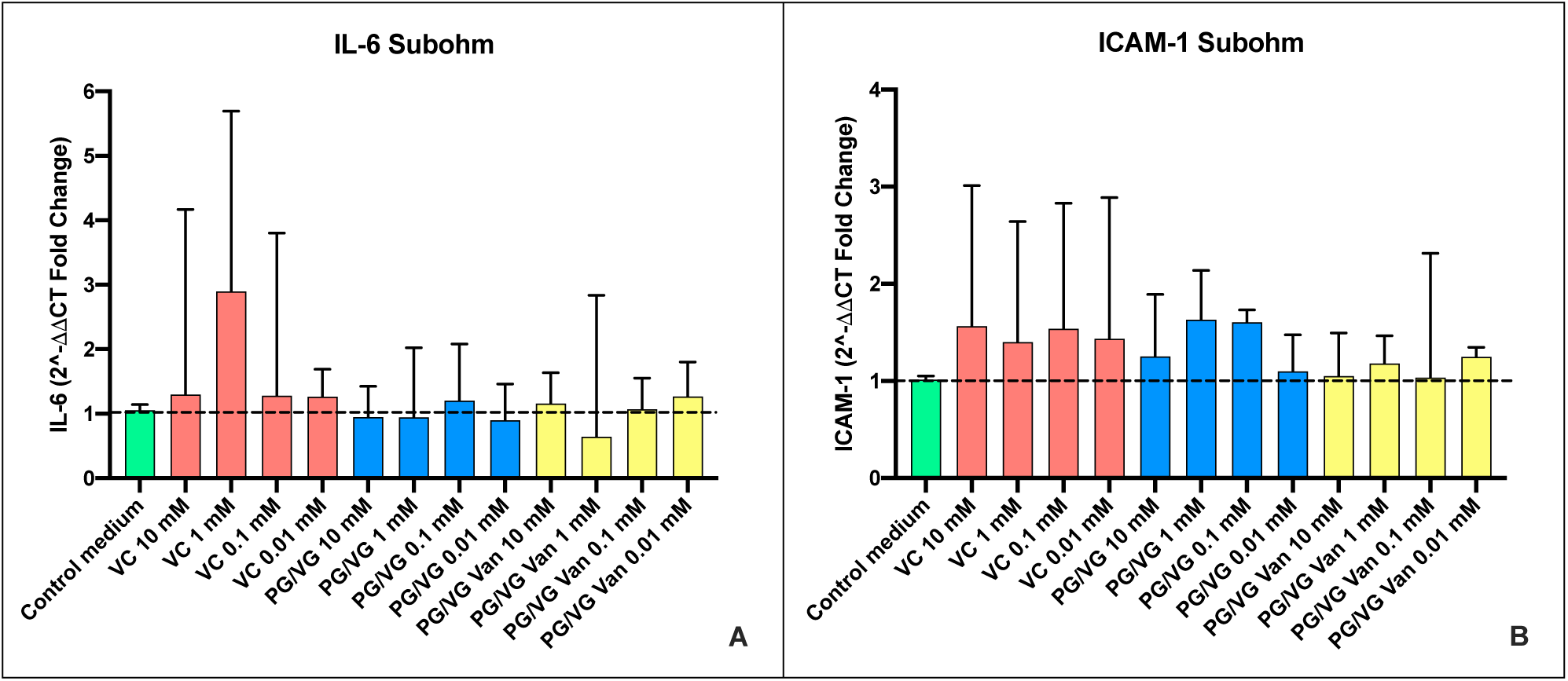
Gene expression of IL-6 (**A**) and ICAM-1 **(B**) in aortic endothelial cells under different treatments produced using the Subohm setting. All data are reported as median (IQR). VC: Vehicle control.

## Discussion

The aim of the present study was the replication of the results by Fetterman et al., 2018 ^16^, with particular regard to investigate the impact of vanillin used in e-liquids on endothelial cell function, evaluating cytotoxicity, oxidative stress, and nitric oxide bioavailability. While the original study provided evidence that vanillin flavoring induces some acute alterations in endothelial functions, our replication efforts revealed different results and covered some methodological gaps mainly concerning the method of exposure.

A key aspect of our replication was the meticulous alignment of our methods with those outlined in the original study and specified personally by the corresponding author. However, despite our efforts to adhere closely to their experimental procedures, we identified several methodological discrepancies that could have contributed to the disparities in our results. Notably, the use of TUNEL assay, which is not typically used for the direct evaluation of cytotoxicity but is commonly employed for detecting apoptotic cell death^20^. Then, TUNEL assay itself does not provide a comprehensive assessment of general cytotoxic effects. For this reason, we performed NRU and MTS assays, which provide information about overall cell damage, regardless of the specific mechanism of cell death ^21^. Moreover, we addressed the main limitation mentioned by the authors: flavor dilution in ethanol rather than propylene glycol (PG) or glycerol (VG), and heating without using an electronic cigarette and a standardized exposure mechanism. We prepared e-liquids by dissolving the vanillin in PG and VG solvents and heating the e-liquid through an electronic cigarette vaped using a standardized vaping machine. We maintained the composition of trapping solution (Ethanol/PBS – 20:80%), as indicated by the corresponding author.

Our results from both NRU and MTS revealed a cytotoxic effect related to the ethanol present in the vehicle control rather than the presence of vanillin in the aqueous extract. It is recognized that ethanol has a cytotoxic effect on cells, especially true at higher concentrations ^22^. This is why we included a vehicle control for each concentration used in the assays, demonstrating that the apparent cytotoxicity of PG/VG and PG/VG with vanillin was mainly due to the presence of ethanol. Beyond Fetterman’s work, there are no other data in the literature indicating cytotoxicity of vanillin, except for cancer cell^23^. In fact, vanillin has been extensively studied as an anti-cancer agent mainly for its anti-proliferative and protective properties against DNA-damaging agents^24, 25^.

Another important property of vanillin is its antioxidant and anti-inflammatory activity ^24, 26^. Vanillin is reported as a potent scavenger of reactive oxygen species (ROS) and an inhibitor of inducible nitric oxide synthase (iNOS)^27^. Assessing oxidative stress, Fetterman and colleagues did not observe a significant increase in ROS after treatment with vanillin. Likewise, our results indicated that vanillin has no pro-oxidant effect on aortic endothelial cells. These results suggest that vanillin, as a component of e-liquids, may not contribute to oxidative stress, which is often a concern with inhalation products.

The bioavailability of nitric oxide (NO) is a critical determinant of endothelial function and play a central role in maintaining vascular homeostasis with its strong vasodilatory, anti-inflammatory, and antioxidant properties^28^. Decreased NO levels have been implicated in endothelial dysfunction, leading to prothrombotic, proinflammatory, and less compliant blood vessel wall^28, 29^. Our investigation into the bioavailability of NO in aortic endothelial cells yielded intriguing results, as we found no significant changes in NO levels in endothelial cells treated with PG/VG vanillin. This contrasts with the reported decrease in NO bioavailability by Fetterman and colleagues, which highlighted discrepancies with existing literature data. Vanillin is well known as a cardioprotective substance^25^. It has been found to stimulate dose-dependent relaxation of isometric tensions during coronary artery contractions induced by different muscle contraction-inducing agents^30^. Furthermore, oral administration of vanillin in mice revealed a cardioprotective effect, reducing cardiac protein oxidation and lipid peroxidation and improving cardiac morphology^31^. Heating vanillin contained in e-liquids could lead to the formation of substances that could impair the endothelial NO balance state, but our results show that acute exposure to PG/VG vanillin under close-to-realistic conditions resulted in no NO reduction in aortic endothelial cells.

Likewise the original study, we evaluated the expression of IL-6 and ICAM-1 genes in aortic endothelial cells under different experimental conditions, specifically focusing on the potential impact of PG/VG and PG/VG vanillin treatments. The outcomes of this study did not reveal any statistically significant variations in IL-6 and ICAM-1 gene expressions when compared to the control, regardless of both experimental settings (Regular or Sub-ohm) or the concentrations used (10 mM, 1 mM, 0.1 mM, and 0.01 mM). The evaluation of IL-6 gene expression under the specified conditions demonstrated no significant differences between the control and treated groups. This observation is consistent across both the Regular and Subohm experimental setups. These findings stand in contrast to the original study by Fetterman et al. (2018)^16^, which reported an overexpression of IL-6 in aortic endothelial cells exposed to 10 mM vanillin. The lack of a significant effect in our replication study suggests that the method of flavor exposure could significantly influence the evaluation of IL-6 expression. The observed discrepancies could be supported by the results of other studies reporting anti-inflammatory properties of vanillin ^24, 25, 32, 33^.

Similar to IL-6, the gene expression of ICAM-1 did not show statistically significant variations under any treatment conditions compared to the control. This result was consistent across all concentrations and both experimental settings. The findings align with the original study by Fetterman et al. (2018) ^16^, which also did not observe any significant effects of vanillin on ICAM-1 gene expression. This consistency reinforces the conclusion that vanillin, at the tested concentrations, does not influence ICAM-1 expression in aortic endothelial cells.

Finally, while commonly used in various scientific applications, ethanol poses potential challenges that require careful consideration, especially when conducting *in vitro* experiments. Cytotoxic ^22^ and pro-oxidant effects ^34^, as well as potential interference with analytical systems are important aspects to consider when evaluating results. It is also essential to consider the ethanol evaporation that researchers must deal with. As observed during our experiments, ethanol evaporation brought negative implications for the reliability and reproducibility of results, with high intra- and inter-laboratory variability.

In conclusion, our replication of the experiments conducted by Fetterman et al., 2018, which focused on the impact of vanillin in e-liquids on endothelial cell function, revealed some disparities in the results compared to the original study. Despite our meticulous efforts to align with the specified experimental procedures, methodological discrepancies - particularly the use of a standardized exposure method closer to real vaping conditions and the choice to test the vehicle control at different concentrations - emerged as critical factors contributing to the observed variations. For the first time, we evaluated the toxicity and cardiovascular effects of vanillin flavor in a context that closely mimics real-world conditions, suggesting that vanillin could be a safer flavoring agent in e-cigarettes, without adverse effects on users’ cardiovascular systems. Further researches are needed to refine our understanding of the potential health effects associated with flavorings in e-liquids. Careful consideration of methodological aspects, particularly the choice of assays and exposure method, is critical for reliable interpretation of results in studies of this nature.

## Highlights

- The evaluation of vanillin flavoring in a context that closely mimics real-use conditions, suggested that vanillin could be a safer ingredient in e-cigarettes, without adverse effects on cardiovascular systems.
- This replication study replicated and clarified the original findings, emphasizing the importance of verifying research results in tobacco harm reduction efforts.
- In vitro studies on the effects of vaping products should be conducted under conditions that closely resemble real-use of these products.
- Replication studies are crucial for validating results, ensuring the robustness of conclusions, identifying errors, improving methodologies, accumulating evidence, and forming scientific consensus. Emphasizing replication strengthens the credibility of science, enhances research practices, and is vital for confirming and validating scientific findings in an era of rapid information dissemination.

## References

1. Agrawal S, Angus K, Arnott D, Ashcroft R, Aveyard P, Barry R, et al. E-cigarettes and harm reduction: An evidence review.

2. Lindson N, Theodoulou A, Ordóñez-Mena JM, Fanshawe TR, Sutton AJ, Livingstone-Banks J, et al. Pharmacological and electronic cigarette interventions for smoking cessation in adults: Component network meta-analyses. Cochrane Database Syst Rev. 2023;9:Cd015226

3. O’Leary R, Polosa R. Tobacco harm reduction in the 21st century. Drugs and Alcohol Today. 2020;20:219–234

4. Patel D, Davis KC, Cox S, Bradfield B, King BA, Shafer P, et al. Reasons for current e-cigarette use among u.S. Adults. Prev Med. 2016;93:14–20

5. DiPiazza J, Caponnetto P, Askin G, Christos P, Maglia MLP, Gautam R, et al. Sensory experiences and cues among e-cigarette users. Harm Reduct J. 2020;17:75

6. Etter JF. Throat hit in users of the electronic cigarette: An exploratory study. Psychol Addict Behav. 2016;30:93–100

7. Caruso M, Emma R, Distefano A, Rust S, Poulas K, Zadjali F, et al. Electronic nicotine delivery systems exhibit reduced bronchial epithelial cells toxicity compared to cigarette: The replica project. Sci Rep. 2021;11:24182

8. Emma R, Fuochi V, Distefano A, Partsinevelos K, Rust S, Zadjali F, et al. Cytotoxicity, mutagenicity and genotoxicity of electronic cigarettes emission aerosols compared to cigarette smoke: The replica project. Sci Rep. 2023;13:17859

9. Caruso M, Emma R, Distefano A, Rust S, Poulas K, Giordano A, et al. Comparative assessment of electronic nicotine delivery systems aerosol and cigarette smoke on endothelial cell migration: The replica project. Drug Test Anal. 2023;15:1164–1174

10. Krüsemann EJZ, Boesveldt S, de Graaf K, Talhout R. An e-liquid flavor wheel: A shared vocabulary based on systematically reviewing e-liquid flavor classifications in literature. Nicotine Tob Res. 2019;21:1310–1319

11. Krüsemann EJZ, Havermans A, Pennings JLA, de Graaf K, Boesveldt S, Talhout R. Comprehensive overview of common e-liquid ingredients and how they can be used to predict an e-liquid’s flavour category. Tob Control. 2021;30:185–191

12. Farsalinos K, Russell C, Polosa R, Poulas K, Lagoumintzis G, Barbouni A. Patterns of flavored e-cigarette use among adult vapers in the USA: An online cross-sectional survey of 69,233 participants. Harm Reduct J. 2023;20:147

13. Dinu V, Kilic A, Wang Q, Ayed C, Fadel A, Harding SE, et al. Policy, toxicology and physicochemical considerations on the inhalation of high concentrations of food flavour. NPJ Sci Food. 2020;4:15

14. Chen T, Wu M, Dong Y, Ren H, Wang M, Kong B, et al. Ovarian toxicity of e-cigarette liquids: Effects of components and high and low nicotine concentration e-cigarette liquid in vitro. Tob Induc Dis. 2023;21:128

15. Effah F, Elzein A, Taiwo B, Baines D, Bailey A, Marczylo T. In vitro high-throughput toxicological assessment of e-cigarette flavors on human bronchial epithelial cells and the potential involvement of trpa1 in cinnamon flavor-induced toxicity. Toxicology. 2023;496:153617

16. Fetterman JL, Weisbrod RM, Feng B, Bastin R, Tuttle ST, Holbrook M, et al. Flavorings in tobacco products induce endothelial cell dysfunction. Arterioscler Thromb Vasc Biol. 2018;38:1607–1615

17. Sinha I, Goel R, Bitzer ZT, Trushin N, Liao J, Sinha R. Evaluating electronic cigarette cytotoxicity and inflammatory responses in vitro. Tob Induc Dis. 2022;20:45

18. Dickinson AJG, Turner SD, Wahl S, Kennedy AE, Wyatt BH, Howton DA. E-liquids and vanillin flavoring disrupts retinoic acid signaling and causes craniofacial defects in xenopus embryos. Dev Biol. 2022;481:14–29

19. ISO International Organization for Standardization. Iso 20768:2018 vapour products — routine analytical vaping machine — definitions and standard conditions. 2018

20. Kyrylkova K, Kyryachenko S, Leid M, Kioussi C. Detection of apoptosis by tunel assay. Odontogenesis: Methods and Protocols. 2012:41–47

21. Riss TL, Moravec RA, Niles AL, Duellman S, Benink HA, Worzella TJ, et al. Cell viability assays. Assay Guidance Manual [Internet]. 2016

22. Nguyen ST, Nguyen HT-L, Truong KD. Comparative cytotoxic effects of methanol, ethanol and dmso on human cancer cell lines. Biomedical Research and Therapy. 2020;7:3855–3859

23. Lirdprapamongkol K, Kramb J-P, Suthiphongchai T, Surarit R, Srisomsap C, Dannhardt G, et al. Vanillin suppresses metastatic potential of human cancer cells through pi3k inhibition and decreases angiogenesis in vivo. Journal of agricultural and food chemistry. 2009;57:3055–3063

24. Arya SS, Rookes JE, Cahill DM, Lenka SK. Vanillin: A review on the therapeutic prospects of a popular flavouring molecule. Advances in Traditional Medicine. 2021;21:1–17

25. Olatunde A, Mohammed A, Ibrahim MA, Tajuddeen N, Shuaibu MN. Vanillin: A food additive with multiple biological activities. European Journal of Medicinal Chemistry Reports. 2022;5:100055

26. Bezerra-Filho CSM, Barboza JN, Souza MTS, Sabry P, Ismail NSM, de Sousa DP. Therapeutic potential of vanillin and its main metabolites to regulate the inflammatory response and oxidative stress. Mini Rev Med Chem. 2019;19:1681–1693

27. Sirangelo I, Sapio L, Ragone A, Naviglio S, Iannuzzi C, Barone D, et al. Vanillin prevents doxorubicin-induced apoptosis and oxidative stress in rat h9c2 cardiomyocytes. Nutrients. 2020;12

28. Tousoulis D, Kampoli AM, Tentolouris C, Papageorgiou N, Stefanadis C. The role of nitric oxide on endothelial function. Curr Vasc Pharmacol. 2012;10:4–18

29. Incalza MA, D’Oria R, Natalicchio A, Perrini S, Laviola L, Giorgino F. Oxidative stress and reactive oxygen species in endothelial dysfunction associated with cardiovascular and metabolic diseases. Vascul Pharmacol. 2018;100:1–19

30. Raffai G, Khang G, Vanhoutte PM. Vanillin and vanillin analogs relax porcine coronary and basilar arteries by inhibiting l-type ca2+ channels. J Pharmacol Exp Ther. 2015;352:14–22

31. Saad HB, Kammoun I, Boudawara O, Hakim A, Amara IB. Preventive effect of vanillin on lipid peroxides and antioxidants in potassium bromate-induced cardiotoxicity in adult mice: Biochemical and histopathological evidences. Journal of Pharmacognosy and Phytochemistry. 2017;6:1379–1383

32. Makni M, Chtourou Y, Fetoui H, Garoui el M, Boudawara T, Zeghal N. Evaluation of the antioxidant, anti-inflammatory and hepatoprotective properties of vanillin in carbon tetrachloride-treated rats. Eur J Pharmacol. 2011;668:133–139

33. Ho K, Yazan LS, Ismail N, Ismail M. Toxicology study of vanillin on rats via oral and intra- peritoneal administration. Food Chem Toxicol. 2011;49:25–30

34. Das SK, Vasudevan DM. Alcohol-induced oxidative stress. Life Sci. 2007;81:177–187

