## Supplemental Materials for "Toxicological evaluation of Vanillin Flavor in E-Liquid Aerosols on Endothelial Cell Function: Findings from the Replica Project"

**Vanillin flavor in e-liquid aerosols does not induce endothelial cell dysfunction: the Replica Project.**

Massimo Caruso

Department of Biomedical and Biotechnological Sciences University of Catania

Via S. Sofia, 97, 95123 Catania, Italy

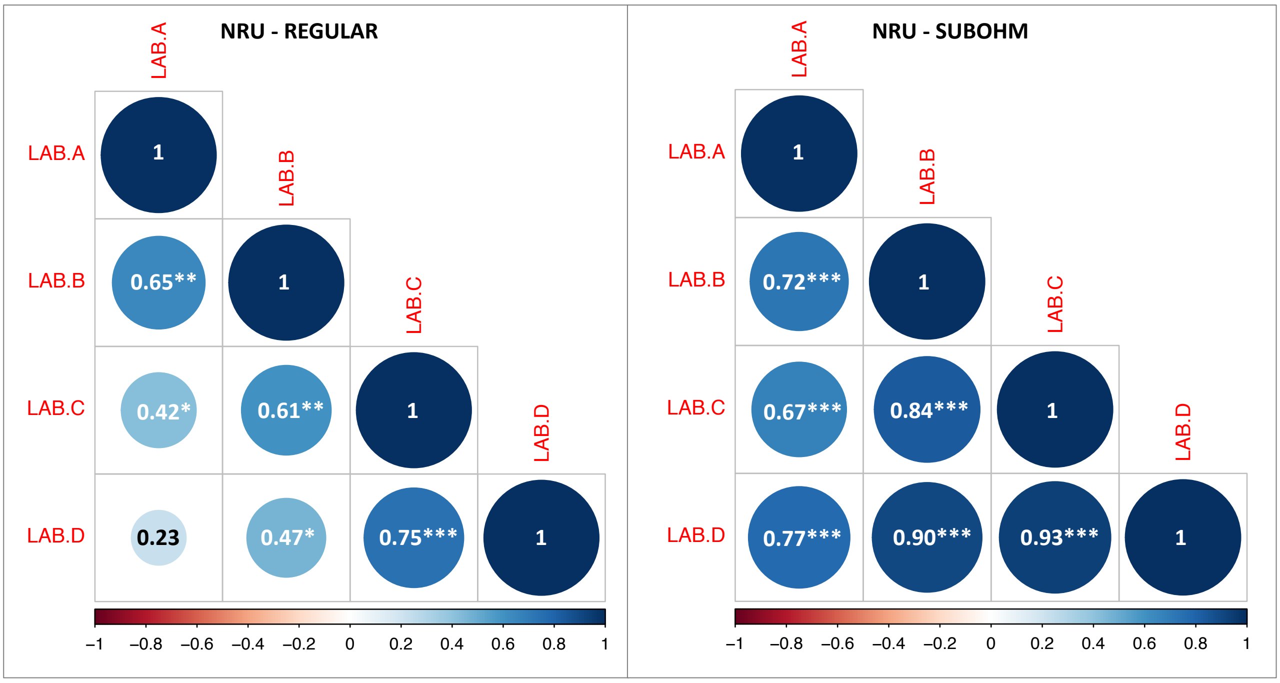


**Figure S1.** Correlograms representing the correlation matrices of NRU data for regular (**A**) and sub-ohm (**B**) settings obtained from each laboratory. Each correlogram shows Spearman's Rank correlation coefficients for all pairs of laboratory data as circles with the corresponding rho value. The color legend on the low side of the correlogram shows the correlation coefficients and the corresponding colors. Positive correlations are displayed in blue and negative correlations in red. The color intensity and the circle size are proportional to the correlation coefficient. Significant correlations were reported as follow: * $p< 0.05$; ** $p< 0.01$; *** $p< 0.001$.

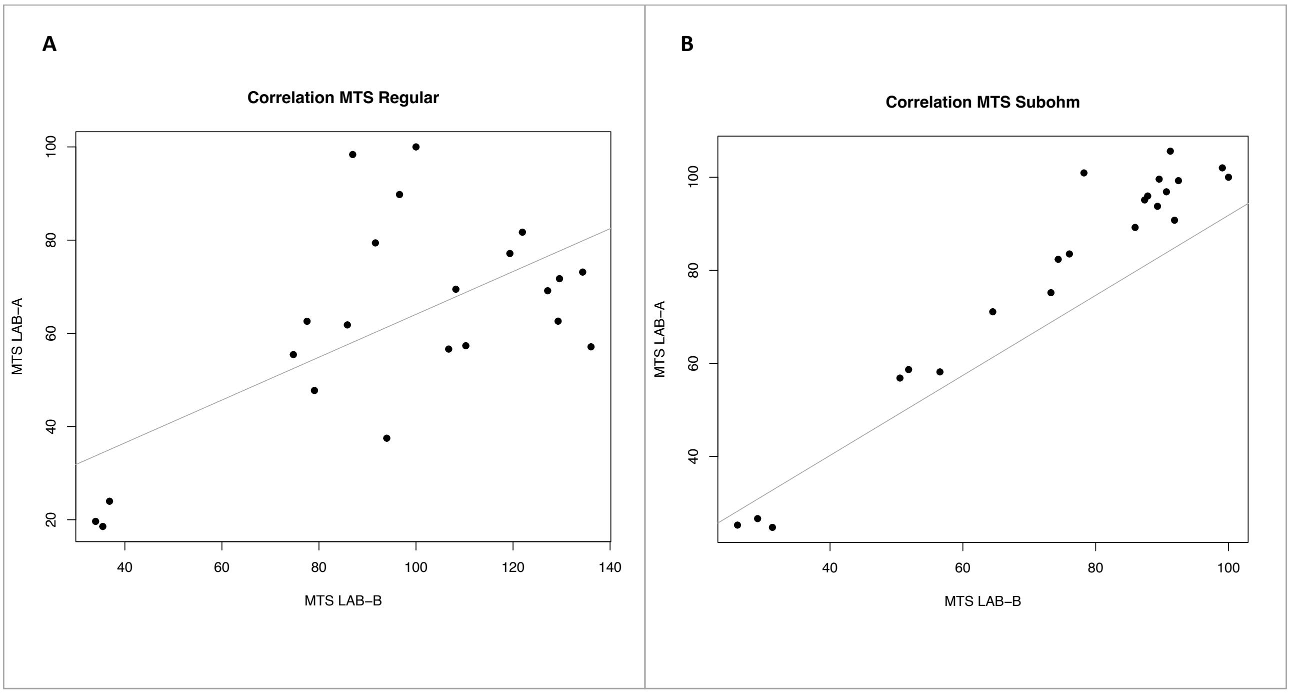


**Figure S2.** Correlation plots representing the correlation of MTS data for regular (**A**) and sub-ohm (**B**) settings obtained from LAB-A and LAB-B. Spearman's Rank correlation coefficients were 0.483 (p= 0.024) for regular setting and 0.896 (p< 0.001) for sub-ohm setting.

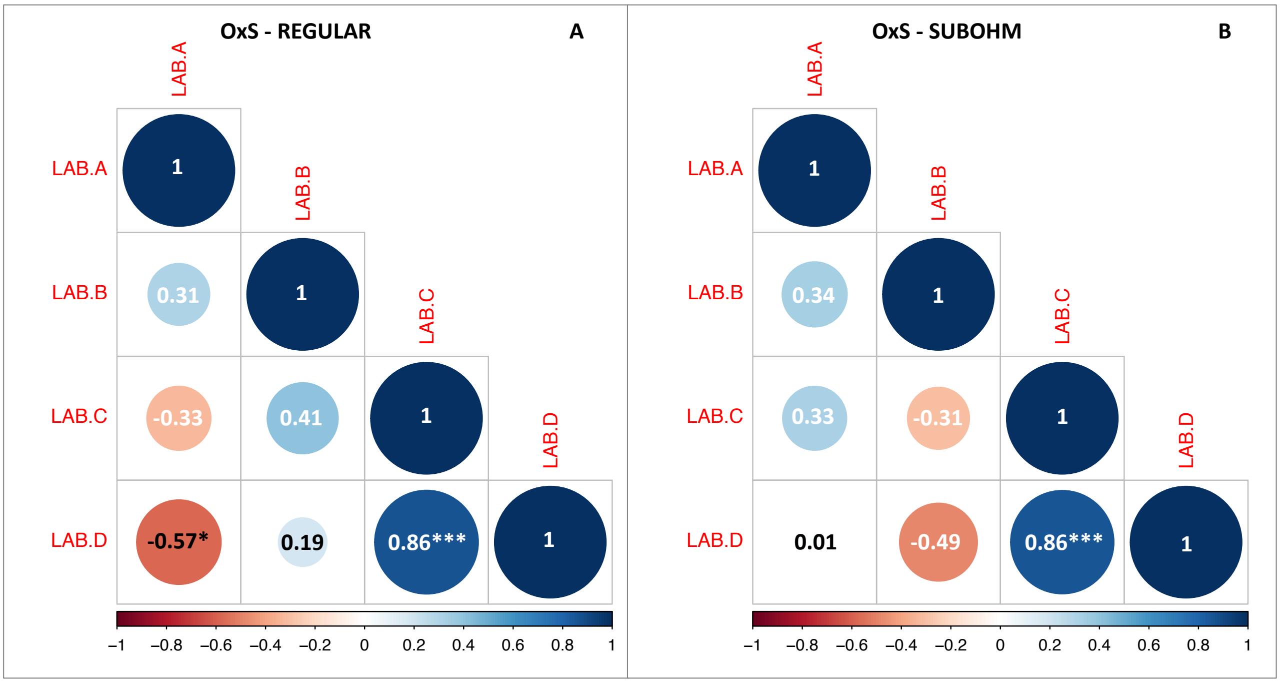


**Figure S3.** Correlograms representing the correlation matrices of oxidative stress (OxS) data for regular (**A**) and sub-ohm (**B**) settings obtained from each laboratory. Each correlogram shows Pearson's correlation coefficients for all pairs of laboratory data as circles with the corresponding R value. The color legend on the low side of the correlogram shows the correlation coefficients and the corresponding colors. Positive correlations are displayed in blue and negative correlations in red. The color intensity and the circle size are proportional to the correlation coefficient. Significant correlations were reported as follow: * p< 0.05; ** p< 0.01; *** p< 0.001.

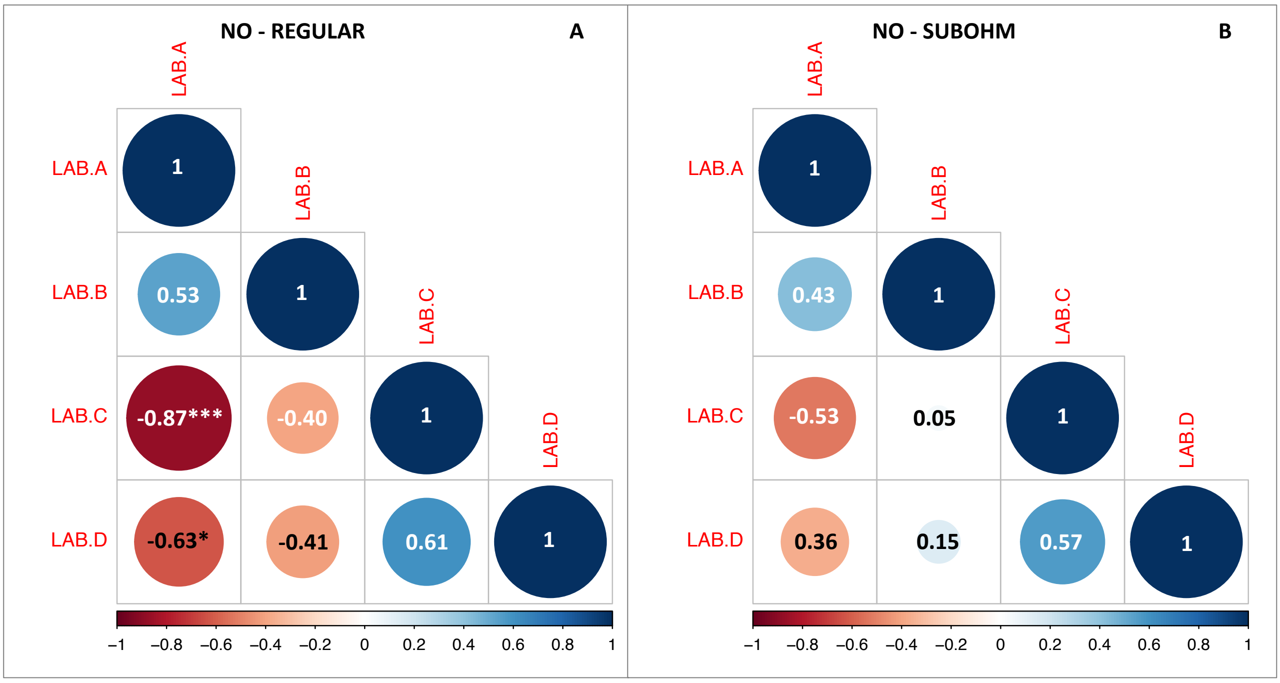


**Figure S4.** Correlograms representing the correlation matrices of NRU data for regular (**A**) and sub-ohm (**B**) settings obtained from each laboratory. Each correlogram shows Pearson's (sub-ohm) and Spearman's Rank (regular) correlation coefficients for all pairs of laboratory data as circles with the corresponding rho/R value. The color legend on the low side of the correlogram shows the correlation coefficients and the corresponding colors. Positive correlations are displayed in blue and negative correlations in red. The color intensity and the circle size are proportional to the correlation coefficient. Significant correlations were reported as follow: * p< 0.05; ** p< 0.01; *** p< 0.001.

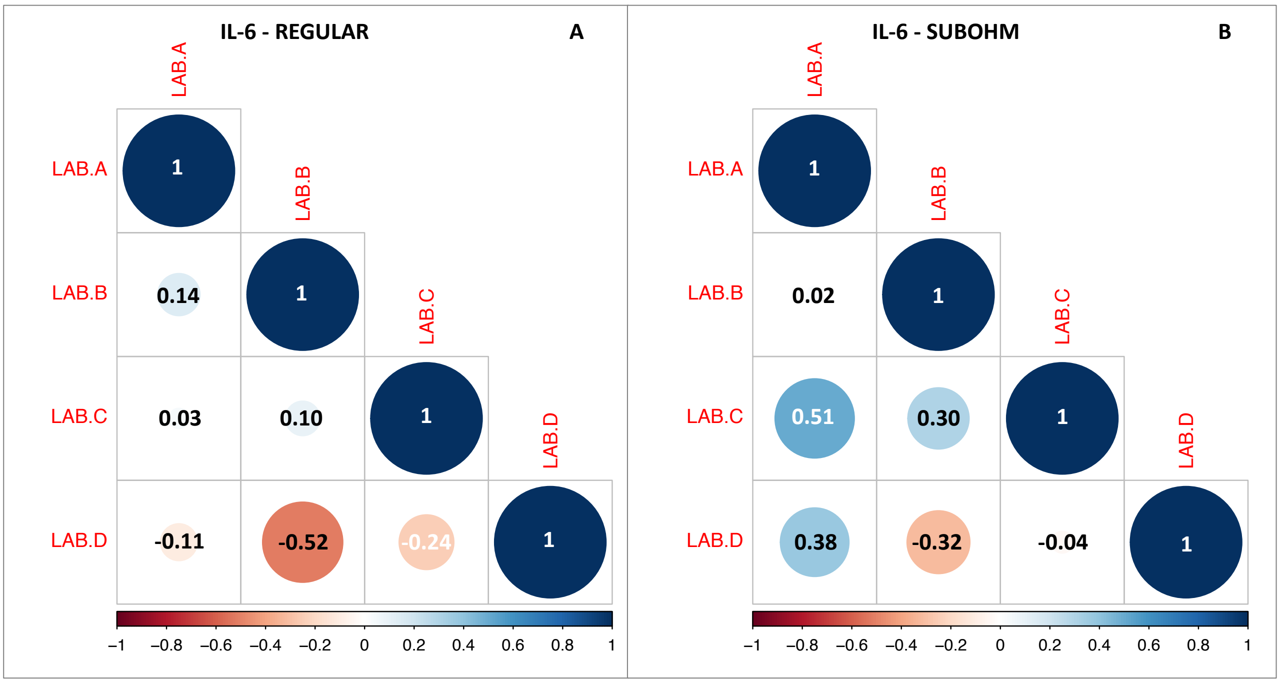


**Figure S5.** Correlograms representing the correlation matrices of IL-6 gene expression data for regular (**A**) and sub-ohm (**B**) settings obtained from each laboratory. Each correlogram shows Spearman's Rank correlation coefficients for all pairs of laboratory data as circles with the corresponding rho value. The color legend on the low side of the correlogram shows the correlation coefficients and the corresponding colors. Positive correlations are displayed in blue and negative correlations in red. The color intensity and the circle size are proportional to the correlation coefficient. No significant correlations were observed.

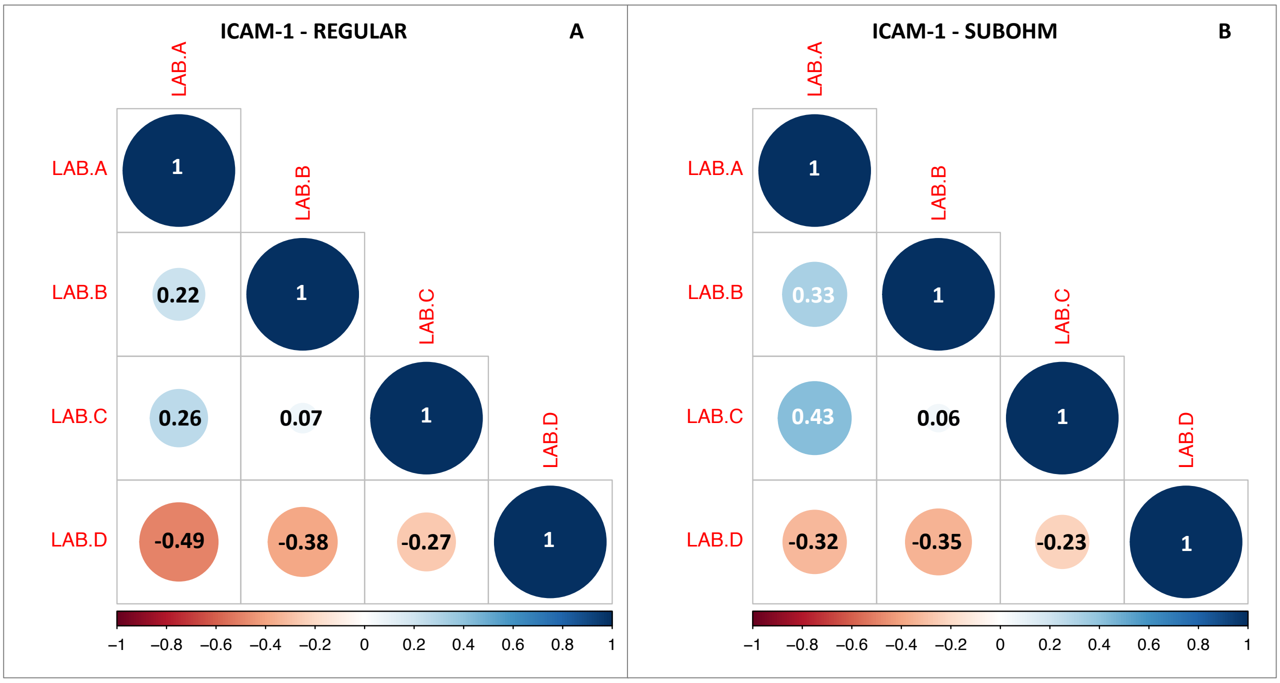


**Figure S6.** Correlograms representing the correlation matrices of ICAM-1 gene expression data for regular (**A**) and sub-ohm (**B**) settings obtained from each laboratory. Each correlogram shows Pearson's correlation coefficients for all pairs of laboratory data as circles with the corresponding R value. The color legend on the low side of the correlogram shows the correlation coefficients and the corresponding colors. Positive correlations are displayed in blue and negative correlations in red. The color intensity and the circle size are proportional to the correlation coefficient. No significant correlations were observed.
